# Ts65Dn synaptic proteome partially recapitulates AD biochemical signature

**DOI:** 10.1101/2023.04.25.538253

**Authors:** Macarena Gómez de Salazar

## Abstract

Down syndrome (DS), caused by human chromosome 21 trisomy, is one of the most common genetic causes of intellectual disabilities. Moreover, approximately 60-80% of individuals with Down Syndrome develop early onset Alzheimer’s disease symptoms and neuropathology characterized by the deposition of senile plaques and neurofibrillary tangles. Spine loss and synaptic deficits are the main alterations present in both pathologies. The synaptic proteins that are altered in both pathologies have also been studied. However, a proteomic study comparing synaptic fractions from DS and AD patients is lacking. To address this aim, we optimized a biochemical subfractioning method to isolate postsynaptic fractions from the cortex of Ts65Dn Down syndrome murine model and the entorhinal cortex from AD subjects’ samples. These samples were used for further proteomic analysis to observe potential alterations in postsynaptic protein expression and protein-protein interaction networks. This study compared the Ts65Dn animal model and entorhinal cortex of AD subjects and revealed a common altered postsynaptic composition of the adult Ts65Dn mouse cortex and entorhinal cortex of AD subjects. These postsynaptic protein alterations might functionally underlie Ts65Dn-associated and AD-associated dendritic deterioration, microtubule organization, glutamatergic transmission, and phosphorylation alterations. They provide insight into the possible common pathological synaptic mechanisms of both diseases and the earliest stages of AD disease progression. Moreover, our proteomic data suggest potential drug targets for modulating the commonly altered postsynaptic mechanisms in DS and AD.

## INTRODUCTION

Trisomy of human chromosome 21 causes Down syndrome (DS), which is one of the most common genetic causes of intellectual disability. A peculiar feature of DS is that approximately 60-80% of individuals with DS will develop early onset Alzheimer’s disease symptoms (Ballard et al., 2016) and neuropathology characterized by the deposition of senile plaques and neurofibrillary tangles (Head et al., 2012). The APP gene is located on chromosome 21 and is suggested to be involved in the relationship between Down Syndrome and Alzheimer’s disease. A proportion of DS individuals will start to accumulate Aβ plaques during their childhood, and almost all DS individuals will have Aβ accumulation in their 20s. By the age of 40, almost all individuals with DS have neurofibrillary tangles (Lemere et al., 1996). However, APP extra copies may explain early onset Alzheimer’s disease features in individuals with DS. Additionally, there are other known processes, such as overexpression of dual-tyrosine kinase 1A (Dyrk1A), which are involved in DS and AD common altered mechanisms. Dyrk1A is present on chromosome 21, and it is consequently overexpressed in DS individuals, with a well-known role in DS pathogenesis (for review see Park et al., 2009). In addition, Dyrk1A protein levels and Dyrk1A mRNA are overexpressed in the postmortem brains of individuals with only AD (Ferrer et al., 2005; Kimura et al., 2007). Dyrk1A aberrantly phosphorylates Tau, leading to neurofibrillary tangle (NFT) accumulation (Ryoo et al., 2007), amyloid precursor protein (APP) (Ryoo et al., 2008) and presenilin 1 (PS1) (Ryu et al., 2010) representing a link between these crucial proteins in AD pathogenesis. Notably, a variety of Dyrk1A inhibitors have been used as therapeutic strategies for the cognitive deficits associated with DS and eventually AD (for review see Smith et al., 2012). For example, in the Ts65Dn mouse model of DS, normalization of Dyrk11A expression recovered memory alterations (Altafaj et al., 2013; García-Cerro et al., 2014). The Ts65Dn mouse model is one of the most well-established murine models of Down Syndrome. Genetically, it consists of the triplication of the Hsa21 syntenic mouse chromosome Mmu16 and recapitulates cognitive and motor alterations observed in individuals with DS (Galdzicki and Siarey, 2003; Altafaj et al., 2013). Moreover, this model has been characterized as an early onset AD murine model (Alldred et al., 2015).

Spine loss and synaptic deficits in DS and AD are among the main alterations present in both pathologies, as described previously in the literature (for review in DS see Cramer and Galdzicki, 2012; for review in AD see Pozueta et al., 2013). In the Ts65Dn mouse model of DS, there is synaptic loss and structural abnormalities in several brain areas (Belichenko et al., 2004). In AD, there is an early and progressive loss of spines, and consequently, synaptic failure provoked by Aβ and tau pathology (for review see Rajmohan and Reddy, 2017). Furthermore, synaptic transmission is altered in both DS and AD. As described in DS individuals (Risser et al., 1997), the Ts65Dn mouse model also presents an imbalance of excitatory/inhibitory neurotransmitter systems (Belichenko et al., 2009). An alteration of GABAergic inhibitory neurotransmission (Fernandez et al., 2009; Best et al., 2012; Contestabile et al., 2017), together with excitatory glutamatergic neurotransmission alterations (Siddiqui et al., 2008; Kleschevnikov, 2012) in Ts65Dn mice, causes this neurotransmitter imbalance. In AD, significantly lower levels of Glutamate and GABA neurotransmitters were measured by HLPC in cortex samples of AD patients than in control subjects (Gueli and Taibi, 2013). More evidence of imbalanced neurotransmission in AD patients with epileptic seizures has also been observed in APP mice with spontaneous nonconvulsive seizure activity in the cortical and hippocampal networks (Palop et al., 2007).

Remarkably, memantine, a reversible N-methyl-d-aspartate glutamate receptor (NMDAR) antagonist, has been used as a treatment for both DS and AD. In the Ts65Dn mouse model, memantine partially rescued the electrophysiological alterations (Scott-McKean and Costa, 2011) and cognitive phenotypic abnormalities (Lockrow et al., 2011). A translation to clinical practice has also shown improvement in the cognitive skills of individuals with DS (Boada et al., 2012). At present, one of the most widely used AD treatments is memantine, which partially alleviates symptoms in AD patients (Reisberg et al., 2003). These results suggest that DS and AD share common molecular mechanisms that may explain synaptic transmission alterations in both pathologies.

The excitatory synapses are structurally formed by a presynaptic terminal, which is a postsynaptic structure with extrasynaptic sides. The postsynaptic fraction contains the postsynaptic density (PSD), which is a molecular scaffold where neurotransmitter receptors such as glutamate receptors are anchored through scaffolding proteins such as PSD95, Shank, and the SAP family (Li et al., 2011a; Naisbitt et al., 1999; Niethammer et al., 1996). Additionally, there are signal transduction proteins, such as kinases and phosphatases, that activate signaling cascades leading to crucial cell processes after receptor activation and Ca2 + influx produced by postsynaptic depolarization. It is possible to biochemically isolate different fractions to study the protein composition of each fraction in detail. Hence, protein-protein interactions within the PSD have been established through extensive mass spectrometry (MS) analyses (For review Shinohara, 2012).

MS analyses have revealed a large degree of molecular complexity in the protein composition that is highly conserved throughout evolution, with approximately more than 1000 proteins in mice and humans within the PSD (Bayés et al., 2017). There is a PSD fraction proteomic study in the Ts65Dn mouse model of Down Syndrome cerebrum (Fernandez et al., 2009). However, this study found only modest changes in the levels of synaptic proteins in the Ts65Dn mouse model. Therefore, we used proteomic analyses to study the postsynaptic and extrasynaptic fractions of the hippocampus of Ts65Dn mice (Gómez de Salazar et al., 2018) to avoid the confounding effect of mixing different brain regions.

Moreover, several proteins involved in synaptic processes have been reported to play a role in both pathologies. Marked synaptic alterations in two actin regulatory proteins that stabilize actin-supporting synaptic plasticity, PAK and the Arp2/3 complex, have been found in adult DS and AD brains (Lauterborn et al., 2020). Another study focused on synaptophysin (SYN) and synaptojanin1 (SYNJ1) levels during DS and AD autopsies. They found that SYN was reduced in DS and DS individuals with AD, whereas SYNJ1 levels (overexpressed in trisomy 21) were significantly higher (Martin et al., 2014).

However, to our knowledge, there are no proteomic studies comparing both DS and AD postsynaptic fraction protein compositions. Therefore, in the present study, we addressed whether DS and AD share disturbances in postsynaptic protein levels that may reflect commonly altered postsynaptic mechanisms in both pathologies. Moreover, through a proteomic study, it was possible to observe postsynaptic protein expression and protein-protein interactions.

To identify these potential molecular synaptic changes, we optimized a biochemical subfractioning method to isolate subsynaptic compartments. Postsynaptic fractions from the cortex of the Ts65Dn Down syndrome murine model and entorhinal cortex from AD subject samples were used for further proteomic analysis. Overall, our results revealed that the Ts65Dn synaptic proteome partially recapitulates the AD biochemical signature, with approximately 40% of commonly regulated proteins involved in functions such as glutamatergic transmission, dendritic spine organization and morphogenesis, cytoskeleton and microtubule binding, synaptic vesicle endocytosis, and phosphorylation. Moreover, these commonly regulated proteins in DS and AD have established protein-protein interaction networks. Modifications of these altered protein networks may represent novel therapeutic targets for DS and AD.

## MATERIALS AND METHODS

### Mice

Ts65Dn mouse colony female B6EiC3Sn a/A-Ts (1716)65Dn (Ts65Dn) and male B6C3F1/J mice were purchased from Jackson Laboratory (Bar Harbor, ME, USA). The mouse colony was housed and bred in the Animal Facilities of Barcelona Biomedical Research Park (PRBB, Barcelona, Spain, EU). All animal procedures met the guidelines of European Community Directive 86/609/EEC and were approved by the Local Ethics Committee. The mice were housed under a 12:12 h light–dark schedule (lights on at 8:00 a.m.) under controlled environmental conditions of humidity (60%) and temperature (22 ± 2°C) with food and water *ad libitum*. Both Ts65Dn and euploid mice were genotyped by qPCR, following the Jackson Laboratories protocol (https://www.jax.org/research-and-faculty/tools/cytogenetic-and-down-syndrome-models-resource/protocols/cytogenic-qpcr-protocol).

### Postmortem human samples

Brain tissue was obtained from the Institute of Neuropathology Brain Bank following the guidelines of Spanish legislation on this matter and the approval of the local ethics committee. Postmortem entorhinal cortex (EC) sections from controls and AD patients were rapidly dissected, frozen on metal plates over dry ice and stored at −80 ◦C until use for biochemical studies. The age from control samples is around 50 years old to prevent the emergence of neurodegeneration features. Post-mortem interval (PMI) are less than ten hours in healthy controls and AD subjects (Figure1). Neuropathologic diagnosis was done following the Braak staging of brain pathology related to AD (Braak and Braak, 1991). Braak stage II samples were selected because they exhibit alterations in EC and are related to the early stages of AD pathology. In this study, one control subject and two AD subjects were studied.

### Subsynaptic Fractionation and Western Blot

The subsynaptic fractionation protocol has been described previously by Gomez de Salazar et al. (2018). This protocol allows the enrichment of extrasynaptic (Extra), postsynaptic (Post), and presynaptic (Pre) fractions from the cortex of 12-month-old mice and the postmortem human entorhinal cortex. In brief, the frozen tissue was thawed in cold buffer A (50 mM Tris-HCl pH 7.4, 0.32 M sucrose, 5 mM EDTA, 1 mM EGTA, 1 μg/ml aprotinin, 1 μg/ml leupeptin, 1/2500 PMSF, 20 μM ZnCl_2_, 50 mM NaF, 1 mM sodium orthovanadate, 2.5 mM sodium pyrophosphate, and a cocktail of protease inhibitors [Complete, Roche]) and mechanically homogenized using a potter. The homogenate was centrifuged for 10 min at 1,400 × g and the supernatant was collected. To increase the yield of protein recovery, this step was repeated three times. The collected supernatants were pooled and centrifuged for 10 min at 700 g. The resulting supernatant was centrifuged again for 15 min at 21,000 g. The resultant pellet containing the crude membrane fraction was kept for further subfractionation as follows: Crude membrane fraction was solubilized in buffer B (0.32 M sucrose, 50 mM Tris-HCl, pH 7.4) and loaded on a discontinuous sucrose step gradient (0.85 M/1.0 M/1.2 M). After centrifugation at 82,500 × g for 2 h, the synaptosomes (Syn) were collected from the 1.0 M/1.2 M interface and diluted in 50 mM Tris-HCl pH 7.4, and centrifuged once more for an additional 30 min at 21,000 × g. Then this pellet was collected and resuspended in synaptosomal resuspension buffer, consisting of 36 mM sucrose 0.1 mM CaCl_2_ + 10 mM NaF + 1 mM sodium orthovanadate, 2 mM EDTA, protease, and phosphatase inhibitor cocktail (Halt^TM^, ThermoFisher), leading to the purified synaptosomal fraction (P2) for further subsynaptic fractionation, as follows. In brief, synaptosomes were solubilized by resuspension in 40 mM Tris-HCl pH 6.0 containing 1% Triton X-100 and incubated at 4°C for 30 min under agitation. Following centrifugation at 40,000 × g for 30 min, the supernatant fraction (corresponding to the extrasynaptic fraction) was precipitated using acetone. The pellet containing the synaptosomal junction was resuspended in 20 mM Tris-HCl buffer pH 8.0 supplemented with 1% Triton X-100. After 30-min of incubation at 4°C, the fractions were separated by centrifugation at 40,000 g for 30 min. The resulting supernatant, corresponding to the enriched presynaptic fraction, was precipitated using acetone. The pellet containing the enriched postsynaptic fraction was resuspended in 50 mM Tris buffer containing 2% SDS, as well as the resulting acetone-precipitated subsynaptic fractions. Only the enriched postsynaptic fraction was preserved and sent to the Mass Spectrometry facility.

### Nano-UPLC Mass Spectrometry System

Mass spectrometry experiments were performed with a pool of two cortices from two different animals for euploid and trisomic conditions, and postmortem human entorhinal cortex samples from one control and two pooled samples from postmortem tissue of patients with AD stage II (Figure1). The samples were subsequently fractionated to obtain subsynaptic fractions from euploid or trisomic mice. Following resuspension and protein quantification, postsynaptic fractions (100–200 μg) were diluted in 1% SDS and digested with the single sequence-specific protease trypsin following the previously described FASP protocol (<u>Wiśniewski et al., 2009</u>). Approximately 45% of each enriched sample was analyzed using an Orbitrap Fusion Lumos with an EASY-Spray nanosource coupled to a nano-UPLC system (EASY-nanoLC 1000 liquid chromatograph) equipped with a 50-cm C18 column (EASY-Spray; 75 μm id, PepMap RSLC C18, 2-μm particles, 45°C). Chromatographic gradients were started at 5% buffer B at a flow rate of 300 nl/min and gradually increased to 22% buffer B in 79 min and to 32% in 11 min. After each analysis, the column was washed for 10 min with 95% buffer B (buffer A:0.1% formic acid in water; buffer B:0.1% formic acid in acetonitrile). The mass spectrometer was operated in data-dependent acquisition mode, with full MS scans over a mass range of *m*/*z* 350–1500 with detection in the Orbitrap (120-K resolution) and auto gain control (AGC) set to 100,000. In each cycle of data-dependent acquisition analysis, following each survey scan, the most intense ions above a threshold ion count of 10,000 were selected for fragmentation with the HCD at a normalized collision energy of 28%. The number of selected precursor ions for fragmentation was determined by the “Top Speed” acquisition algorithm (maximum cycle time of 3 s), and a dynamic exclusion of 60 s was set. Fragment ion spectra were acquired in an ion trap with an AGC of 10,000 and maximum injection time of 200 ms.

### Raw Data Processing and Data Analysis

The acquired data were analyzed using the Proteome Discoverer software suite (v2.0, Thermo Fisher Scientific), and the Mascot search engine (v2.5.1, Matrix Science) was used for peptide identification. Data were searched against a *Mus musculus or Homo Sapiens* protein database derived from SwissProt, which included the most common contaminants. A precursor ion mass tolerance of 7 ppm at the MS1 level was used, and up to three missed cleavages of trypsin were allowed. The fragment ion mass tolerance was set to 0.5 Da oxidation of methionine. The identified peptides were filtered using a 5%FDR.

### Bioinformatic Analysis of Proteomic Data

The proteomic signatures of the experimental groups were represented using Venn diagrams. These diagrams were created using VENNY 2.1 freeware (<u>Oliveros et al., 2007–</u> <u>2015</u>) (http://bioinfogp.cnb.csic.es/tools/venny/). These graphical interfaces allowed the representation of proteins that overlapped among groups, together with the visualization of differentially expressed proteins in the different groups/fractions. Additionally, GraphPad Prism 8 was used to generate heat maps. These values represent the logarithmic scale of the protein expression ratio. When this ratio is 0.75-1.25, we considered that proteins did not change between the control condition and pathology. When the ratio is less than 0.75 proteins are considered to be downregulated, and when it is more than 1.25 proteins are considered to be upregulated.

To functionally classify the differentially expressed proteins from the different groups (genotypes), the data were submitted to the EnrichR software (Kuleshov et al., 2016). This software, based on gene ontology (GO), was used to group differentially expressed proteins into biological processes, molecular functions, and cellular components.

Analysis of the proteome signatures was performed using the identification of proteins in the biological samples as inclusion criteria. Regarding the latter, proteins detected in the postsynaptic fractions were classified based on the relative TS:EU and Control:AD ratios. Ratios < 0.75 or >1.5 were considered as downregulated and upregulated protein levels, respectively. Finally, interaction network analysis was performed using the Search Tool for the Retrieval of Interacting Genes/Proteins database (STRING; http://www.string-db.org/). This bioinformatics tool allowed the generation of protein–protein interaction networks.

### Statistical Analysis

The adjusted p-value using the Benjamini-Hochberg method for correction of multiple hypotheses testing for the enrichment of proteins in the different gene ontology groups was considered.

## RESULTS

### Postsynaptic Proteomic Signature of the Ts65Dn Mice and Postmortem AD samples

Proteomic studies have shown that synaptic mechanisms are affected individually in DS and AD models. However, there is not much evidence showing the specific changes in the protein composition of the postsynaptic compartments that may impact synaptic function in both AD and DS. Specifically, identifying common alterations in both pathologies could provide a novel perspective on the mechanisms that lead to the development of early onset AD in individuals with DS. To address this issue, we performed a subsynaptic fractionation protocol to obtain extrasynaptic-, presynaptic-, and postsynaptic-enriched fractions, both from euploid, trisomic adult mouse cortex and human postmortem entorhinal cortex (EC) from control and AD patients. Postmortem EC from human subjects was selected for our study because EC in the medial temporal lobe is already affected in the early stages of AD (Braak and Braak, 1991). The cortical subsynaptic fractionation protocol was previously validated by our group by western blotting using specific subsynaptic compartment markers (Gomez de Salazar et al., 2018).

Postsynaptic fractions were collected from the four groups for mass spectrometry analysis (Figure 1). Postsynaptic fractions were subjected to trypsin digestion, and mass spectrometry was performed. Data analysis was based on a comparison of the effects of genotype on the postsynaptic proteomic signature. Proteomic analysis allowed the identification of 5916 proteins (M. musculus protein database and H. Sapiens protein database, SwissProt) from the four groups. A distribution analysis of these proteins showed that 1491/1205 proteins were present in the postsynaptic fractions of the cortex from euploid/trisomic mice, whereas 1660/1560 proteins were present in the postsynaptic fractions of the entorhinal cortex from control and AD subjects, respectively (Figure 2A and Supplementary Table 1). The proteomic profile showed the presence of 600 conserved proteins that are present in all the studied groups: Euploid mice, Trisomic mice, Control human, and AD Human. Further in silico analyses were performed to analyze potential functional proteomic changes in the group of conserved proteins across species and genotypes to understand the common alterations in postsynaptic protein levels and functions in DS and AD pathologies.

**Figure 1.**
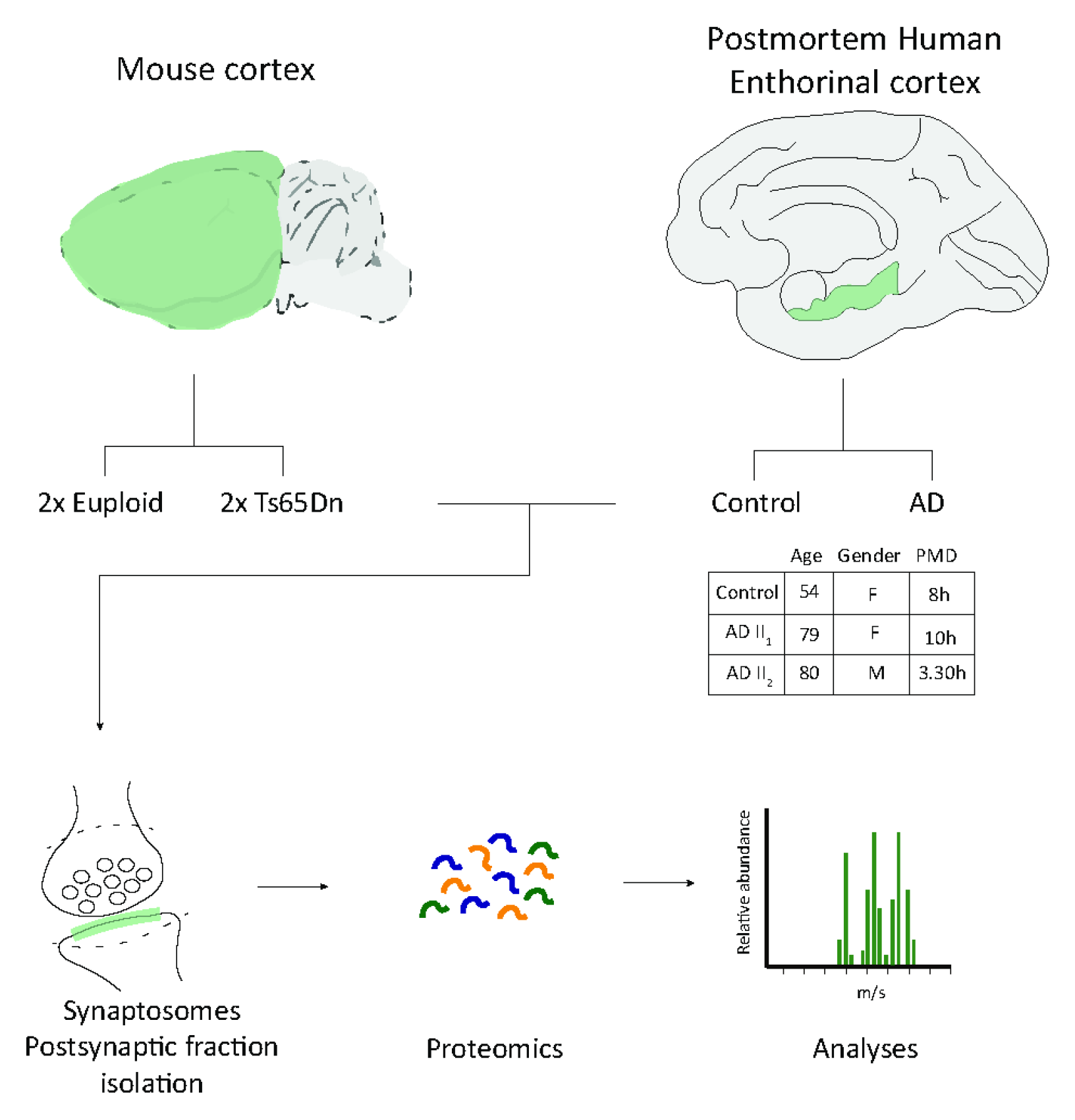
Proteomic study workflow. Cortex from euploid (n=2) and trisomic mice (n=2) and entorhinal cortex postmortem tissue from controls (n=1) and AD subjects (n=2). Subsynaptic fractionation and proteomic analysis of postsynaptic fractions from Ts65Dn mouse cortex and postmortem entorhinal cortex of AD subjects.

**Figure 2.**
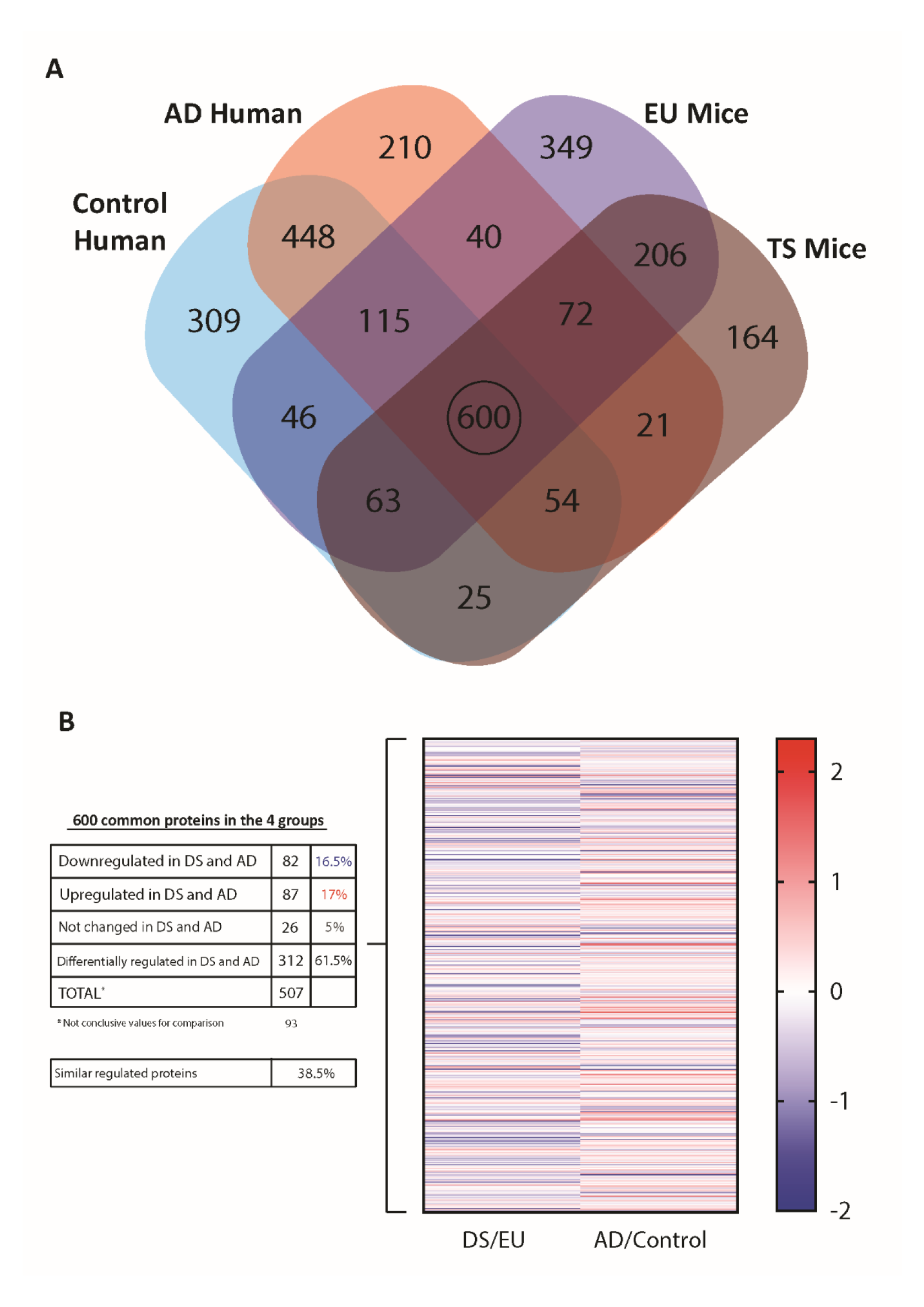
Common proteins expressed in postsynaptic fraction of cortex from euploid and trisomic mice and entorhinal cortex postmortem tissue from controls and AD subjects. A) Venn’s diagram representing total identified proteins overlap between postsynaptic fractions from the 4 genotypes (600 proteins). (B) Table representing the quantity of proteins that are affected or not changed between DS and AD pathologies. Heatmap diagram representing ratios (0.75-1.25) from trisomic vs. euploid mice (DS/EU) and ratios from AD subjects vs. control subjects (AD/Control). Lines in red: up-regulated proteins (<1.25); Lines in blue: down-regulated proteins (>0.75).

### Altered protein expression in the Ts65Dn Mice Cortex and Postmortem AD entorhinal cortex samples

Next, the 600 conserved proteins that were present in all studied groups were analyzed by comparing protein expression values by obtaining ratios from trisomic vs. euploid mice (DS/EU) and ratios from AD subjects vs. control subjects (AD/Control). The values for 93 proteins were not conclusive enough to obtain ratios. Hence, the DS/EU and AD/Control ratios of 507 proteins were obtained for further studies. These ratios are represented in a heatmap diagram (Figure 2B, right; Supplementary Table 2). The number and percentage of proteins that were similarly or differently regulated in DS and AD are organized in a table (Figure 2B, left). A subset of 312 proteins (61.5% of total proteins) was differentially regulated in DS and AD. Remarkably, 195 proteins (38.5% of the total proteins) were regulated in the same direction. A subset of 26 proteins (5% of the total proteins) was not changed in either DS or AD. In contrast, a subset of 82 proteins was downregulated in both DS and AD samples (16.5% of total proteins). Additionally, a subset of 87 proteins was upregulated in both DS and AD samples (17% of total proteins). To further understand the common synaptic alterations in both pathologies, functional annotation analyses were performed on both subsets of proteins that were similarly downregulated or upregulated in DS and AD.

### Gene Ontology Enrichment Analysis of postsynaptic Ts65Dn Mice Cortex and Postmortem AD entorhinal cortex samples

The subsets of 82 downregulated or 87 upregulated proteins in DS and AD were subjected to GO-term-based protein enrichment analysis using EnrichR software. For these functional annotation studies, the groups of downregulated or upregulated proteins were studied separately. Statistical enrichment analysis of “biological processes,” “cellular compartment,” and “molecular functions” revealed the presence of significant differences between upregulated and downregulated proteins (Figures 3A-C, 4A-C).

**Figure 3.**
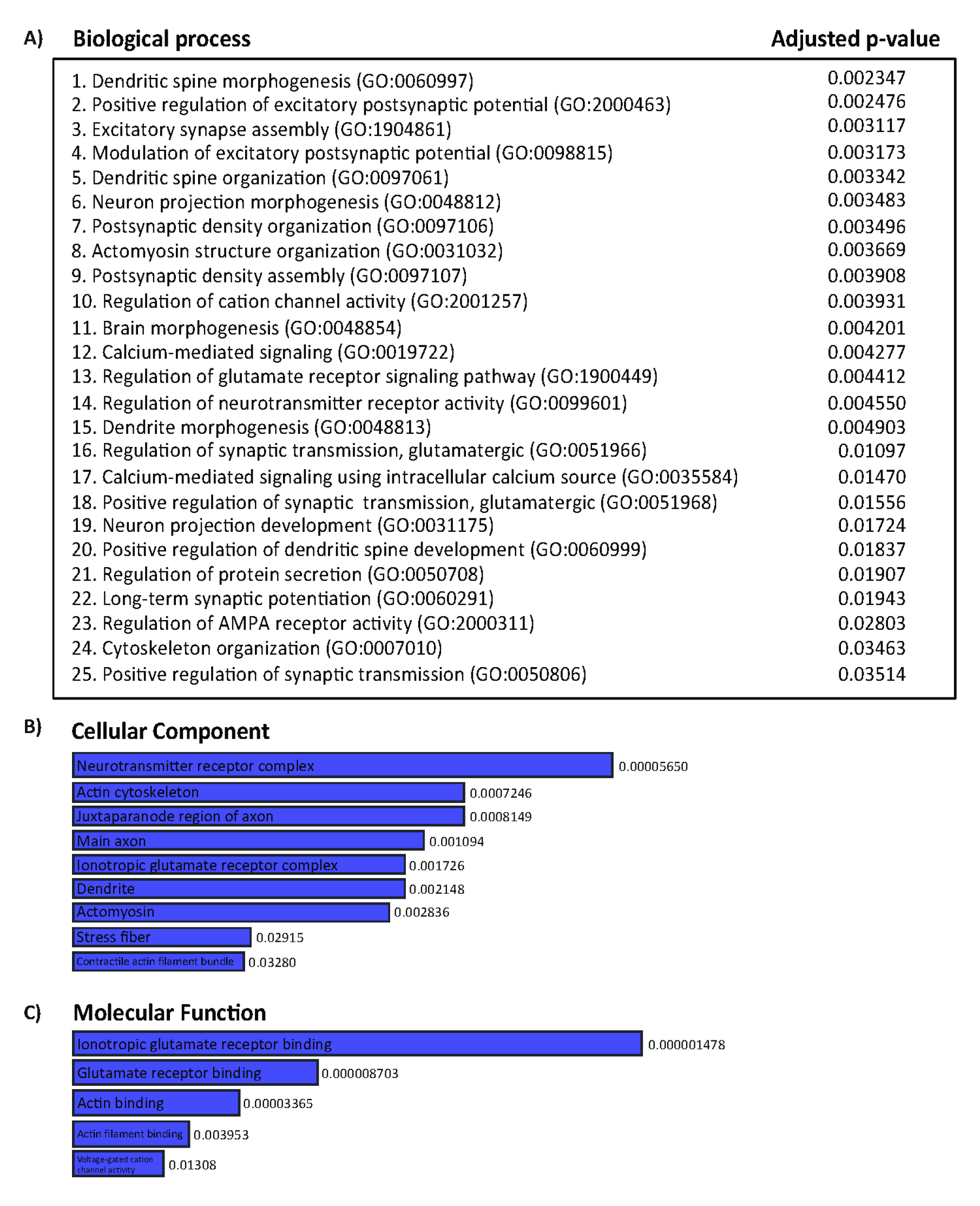
Gene ontology analysis of common proteins that are downregulated in postsynaptic fractions from trisomic cortex and entorhinal cortex postmortem tissue from AD subject. Functional annotation analysis of proteins enriched in both postsynaptic fractions from DS and AD. (A) Table listing significantly enriched protein groups related with “biological process”. (B) Bar graphs representing significantly enriched GO terms, based on “cellular compartment” criteria. (C) Bar graphs representing significantly enriched GO terms, based on “molecular function” criteria, in the postsynaptic fraction of Down Syndrome (D) and AD samples.

The downregulated cluster of 82 protein analyses (Figures 3A-C), principally revealed that functions such as dendrite, dendritic spine morphogenesis and organization, postsynaptic density organization and assembly, excitatory synapse assembly, cytoskeleton organization, actin cytoskeleton, actin binding, and actomyosin structure organization were significantly enriched. These downregulated functions suggest that the organization and assembly of dendrites and dendritic spines are similarly altered in DS and AD. Moreover, functions such as positive regulation of excitatory postsynaptic potential, modulation of excitatory postsynaptic potential, regulation of glutamate receptor signaling pathway, regulation of glutamatergic synaptic transmission, neurotransmitter receptor complex, ionotropic glutamate receptor complex, ionotropic glutamate receptor binding, glutamate receptor binding, long-term potentiation (LTP), and regulation of AMPA receptor activity were also significantly enriched within the downregulated cluster of proteins, suggesting that excitatory glutamatergic transmission is also altered in the cortex of Ts65Dn mice and the EC of AD subjects.

The upregulated cluster of 87 proteins analyses (Figures 4A-C), revealed that functions such as cytoskeleton, microtubule, microtubule and tubule binding, axon, axon development, and neuron projection morphogenesis development were significantly enriched. These upregulated functions suggest that the development of neuronal projections may be similarly altered in DS and AD. Additionally, these functional analyses revealed that protein kinase activity, protein ser/threo kinase activity, and phosphorylation were enriched in the upregulated protein cluster, suggesting an alteration in protein kinase function and phosphorylation. In addition, vesicle functions, such as synaptic vesicle endocytosis, clathrin endocytosis, and clathrin-sculpted GABA transport vesicles, were enriched in this protein cluster.

**Figure 4.**
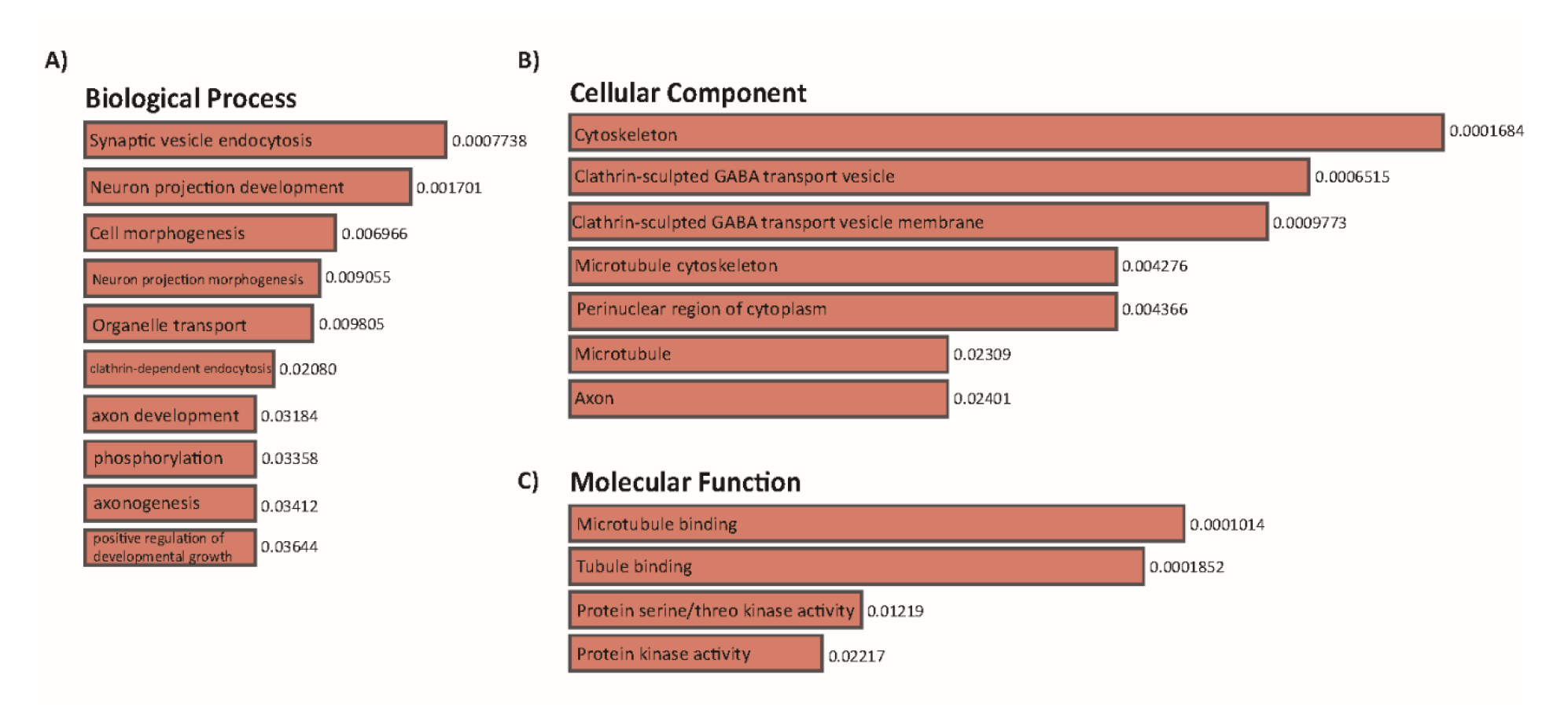
Gene ontology analysis of common proteins that are downregulated in postsynaptic fractions from trisomic cortex and entorhinal cortex postmortem tissue from AD subject. Functional annotation analysis of proteins enriched in both postsynaptic fractions from DS and AD. (A) Bar graphs representing significantly enriched GO terms, based on “biological process” criteria. (B) Bar graphs representing significantly enriched GO terms, based on “cellular compartment” criteria. (C) Bar graphs representing significantly enriched GO terms, based on “molecular function” criteria, in the postsynaptic fraction of Down Syndrome (D) and AD samples.

Overall, GO-term analysis suggested that the postsynaptic proteome of the trisomic mouse cortex and entorhinal cortex of AD subjects is altered when compared to controls, in terms of both cellular compartments and functional outcomes. Remarkably, both postsynaptic proteomes share features that are simultaneously altered in DS and AD and may underlie some of the pathological synaptic alterations present in both conditions.

### Protein-protein interaction networks in postsynaptic Ts65Dn Mice Cortex and Postmortem AD entorhinal cortex samples

To better understand the links between these proteins, potential protein–protein interaction networks between downregulated and upregulated clusters of proteins in DS and AD were built using String software. Interactome analyses of the downregulated cluster of 82 proteins revealed a significant interaction between this group of proteins, as shown by the elevated number of edges (predicted number of edges:80 vs. expected number of edges:43) and PPI enrichment (p-value =2.62e-07), indicating significant interactions within the PPI network. STRING analysis identified in the center of the network DLG4, which in humans refers to PSD-95, one of the most studied members of the MAGUK family of PDZ domain-containing proteins. PSD-95 distinctively interacts with other proteins from the postsynaptic scaffold, such as SHANK family proteins, HOMER2, glutamate receptors (mGluR5), and kinases from the CAMKII family (Figure 5).

**Figure 5.**
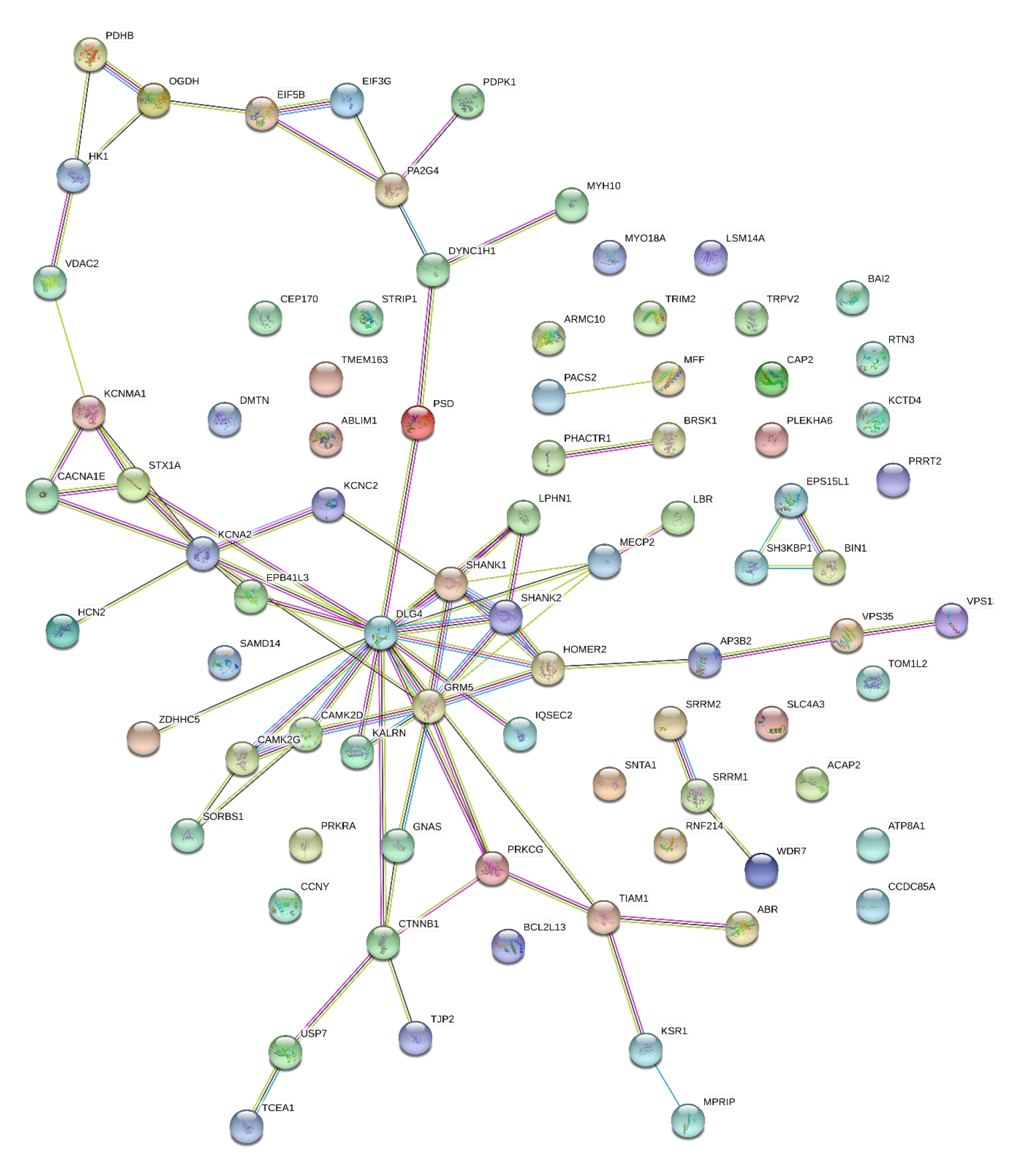
Protein–protein interaction (PPI) network of protein downregulated in both trisomic postsynaptic fraction and AD postsynaptic fraction. Node color annotation: red, kinase activity; blue, postsynaptic density-associated kinase; green, participation in long-term potentiation; and yellow, present in the glutamatergic synapse; link color annotation: known interactions (light blue: curated databases; pink: experimentally determined), predicted interactions (green: gene neighborhood; red: gene fusions; blue: gene co-occurrence), and others (light green: text mining; black: coexpression; purple: protein homology)

Interactome analyses of the upregulated cluster of 87 proteins revealed a significant interaction between this group of proteins, as shown by the elevated number of edges (predicted number of edges:99 vs. expected number of edges:42) and PPI enrichment (p-value = 5.02e-14), indicating significant interactions within the PPI network. STRING analysis identified proteins such as synaptotagmin 1 (SYT1), PACSIN1, ITSN1, NCAM1, NCAM2, GAD1, and GABBR1 that distinctively interact with each other and with other proteins (Figure 6).

**Figure 6.**
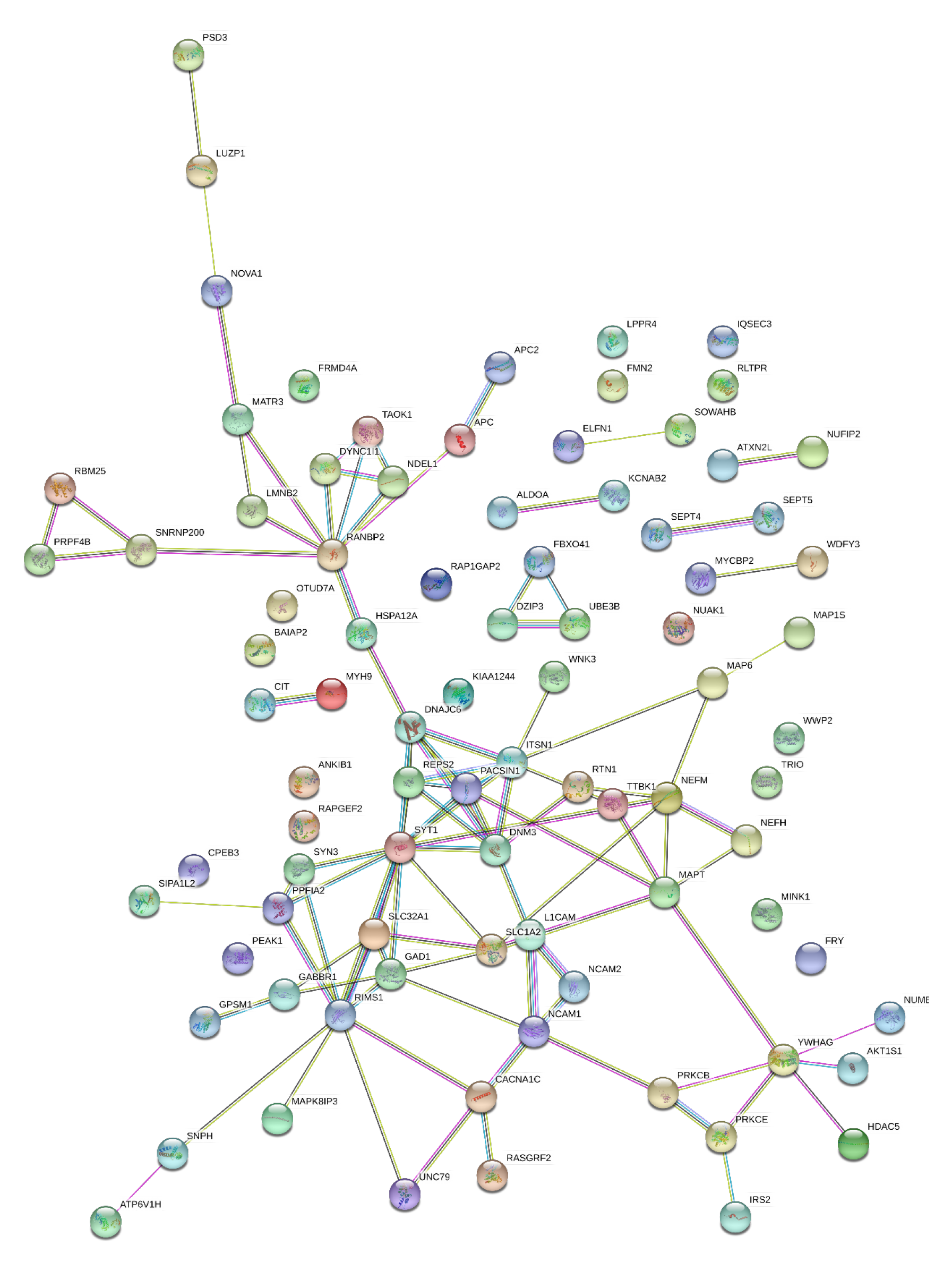
Protein–protein interaction (PPI) network of protein upregulated in both trisomic postsynaptic fraction and AD postsynaptic fraction. Node color annotation: red, kinase activity; blue, postsynaptic density-associated kinase; green, participation in long-term potentiation; and yellow, present in the glutamatergic synapse; link color annotation: known interactions (light blue: curated databases; pink: experimentally determined), predicted interactions (green: gene neighborhood; red: gene fusions; blue: gene co-occurrence), and others (light green: text mining; black: coexpression; purple: protein homology)

### Altered kinase expression in the Ts65Dn Mice Cortex and Postmortem AD entorhinal cortex samples

Subsequently, 53 common conserved kinases that were present in all the studied groups were analyzed by comparing protein expression values by obtaining ratios from trisomic vs. euploid mice (DS/EU) and ratios from AD subjects vs. control subjects (AD/Control).. The number and percentage of kinases that were similarly or differently regulated in DS and AD are organized in a table and represented in a heatmap (Figure 7, Supplementary Table 3). A subset of 37 kinases (58.5% of total kinases) was differentially regulated in DS and AD. Remarkably, 21 kinases (39.6% of the total proteins) were regulated in the same direction. A subset of 26 kinases (18.9% of total proteins) did not change in either DS or AD. In contrast, a subset of 82 proteins was downregulated in both DS and AD samples (16.5% of total proteins). Additionally, a subset of 87 proteins was upregulated in both DS and AD samples (17% of total proteins). To further understand the common synaptic alterations in both pathologies, functional annotation analyses were performed on both subsets of proteins that were similarly downregulated or upregulated in DS and AD.

**Figure 7.**
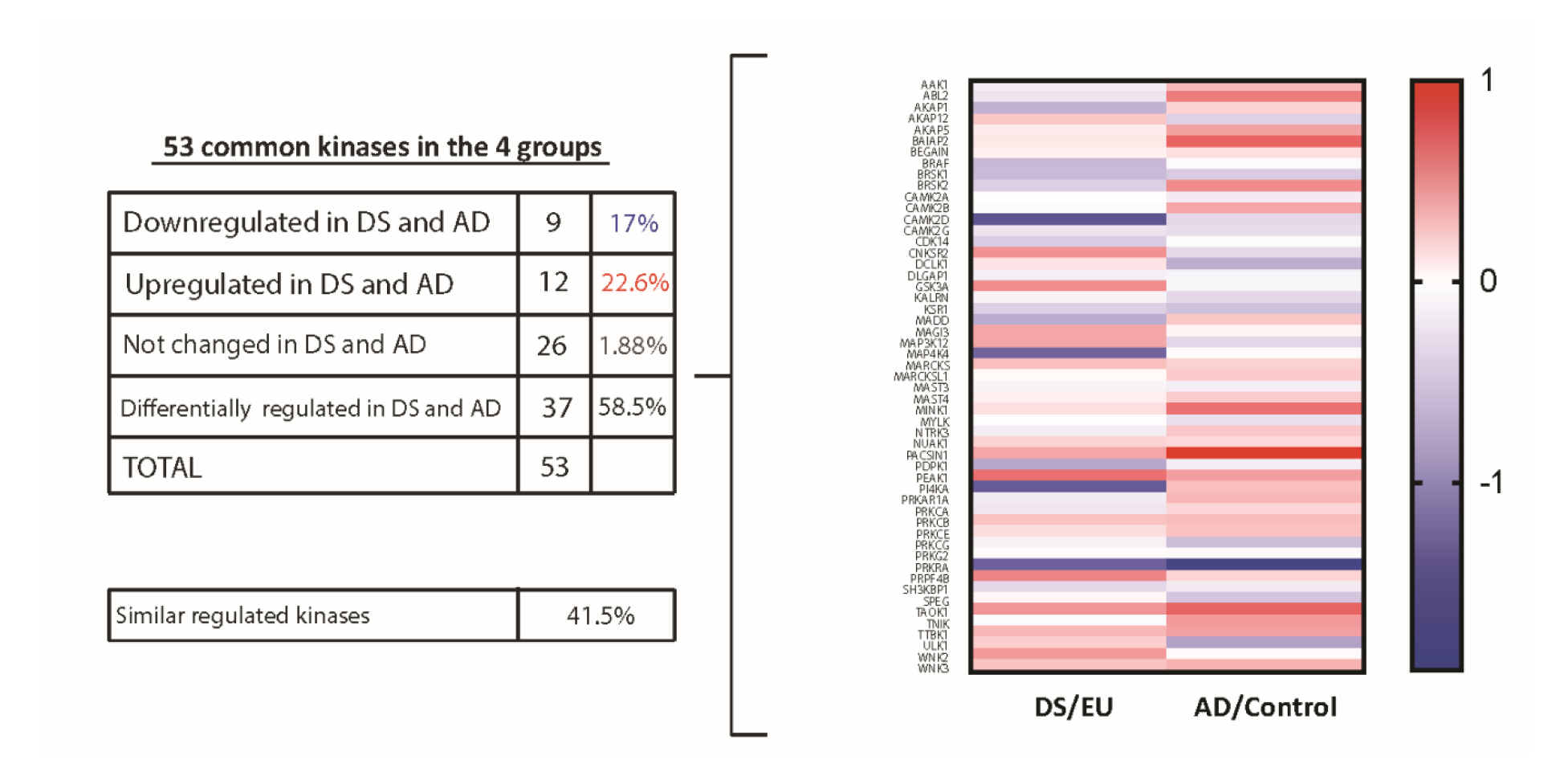
Common kinases expressed in postsynaptic fraction of cortex from euploid and trisomic mice and entorhinal cortex postmortem tissue from controls and AD subjects. Table representing the quantity of proteins that are affected or not changed between DS and AD pathologies. Heatmap diagram representing ratios (0.75-1.25) from trisomic vs. euploid mice (DS/EU) and ratios from AD subjects vs. control subjects (AD/Control). Lines in red: up-regulated kinases (<1.25); Lines in blue: down-regulated kinases (>0.75).

## DISCUSSION

In this study, we optimized a biochemical subfractioning method to isolate postsynaptic fractions from the cortex of Ts65Dn Down syndrome murine model and the entorhinal cortex from AD subjects’ samples. These samples were used for further proteomic analysis to observe alterations in postsynaptic protein expression and protein-protein interaction networks.

Overall, our results revealed that Ts65Dn synaptic proteome partially recapitulates AD biochemical signature.

Approximately 6000 proteins from mice and human postsynaptic samples were detected in the four groups. In each of the postsynaptic samples from the cortex of euploid/trisomic mice, approximately 1500 proteins were found in each of the postsynaptic samples from the cortex of euploid/trisomic mice and the entorhinal cortex from control/AD subjects. This number of proteins is in line with previous studies that demonstrated that mouse and human postsynaptic density in cortical samples comprised approximately 1500 proteins (Bayès et al., 2012). Our proteomic analyses indicated the presence of 600 conserved proteins that are present in all the studied groups: Euploid mice, Trisomic mice, Control human, and AD Human. Approximately 40% of proteins found in all groups (murine and postmortem human tissue samples) were regulated in the same direction. The analyses revealed that 16.5% of proteins were downregulated and 17% were upregulated in DS and AD versus controls, while 5% did not change. In contrast, approximately 60% of the proteins were regulated differently between DS and AD.

Further GO-term-based protein enrichment analysis indicated that functions such as dendritic function, postsynaptic density organization, cytoskeleton organization, calcium-mediated signaling, and excitatory glutamatergic transmission were affected in the downregulated protein clusters in DS and AD versus controls. These findings agree with previous results from our lab showing that the hippocampal postsynaptic fraction of trisomic mice showed a relative decrease of proteins related with “dendritic function” GO term (Gómez de Salazar et al., 2018). These results are also in agreement with previous studies showing a reduction in dendritic branching and spine density in Ts65Dn mice (Belichenko et al., 2007) and dendritic deterioration reported in DS fetal brains (Weitzdoerfer et al., 2001). There is also evidence of dendritic abnormalities in AD (for review see Cochran et al., 2014).

Moreover, previous studies have described that PSD95 levels are dismissed in the Ts65Dn mouse model (Alldred et al., 2015) and in the human temporal cortex of AD individuals (Gylys et al., 2004) in line with the downregulation of this crucial postsynaptic protein that we have found in our proteomic study. Another function downregulated in the postsynaptic protein cluster in DS and AD is calcium-mediated signaling. Previous studies have shown that calcium signaling is dysregulated in the TgDyrk1A mouse cerebellum (Altafaj et al., 2013) and disruption of intracellular Ca2+ homeostasis in AD (for review see Wang et al., 2017a) in agreement with our proteomic data.

Moreover, our results indicate a common excitatory glutamatergic transmission alteration in the cortex of Ts65Dn mice and the EC of AD subjects. As described in this study, oral administration of memantine to Ts65Dn mice improved spatial learning and reduced cortical and hippocampal AβPP levels in aged TS mice (Rueda et al., 2010), indicating a glutamatergic transmission alteration that may share common mechanisms in Ts65Dn mice as a model of Down Syndrome and early onset Alzheimer’s disease. In particular, crucial functions for synaptic plasticity, such as long-term potentiation (LTP) and regulation of AMPA receptor activity, were enriched in the downregulated cluster of common postsynaptic proteins in DS and AD from our samples. For example, hippocampal LTP has been reported to be reduced in Ts65Dn mice (Siarey et al., 1997; Kleschevnikov et al., 2004) and in the perirhinal cortex (Roncacé et al.,2017). Additionally, it is well established that LTP is impaired in different transgenic AD models (for review see McGowan et al., 2016). The accumulation of soluble Aβ oligomers specifically blocks cortical and hippocampal LTP (Shankar et al, 2008). In AD patients, LTP-like cortical plasticity has been reported to be also altered in primary motor cortex (Koch et al, 2012). Biochemical analysis of AMPARs has revealed a reduction in AMPAR surface expression in the hippocampus of Tc1 mice (Morice et al., 2008). Another mechanism that may explain this reduction in AMPAR surface expression is that Sorting Nexin (SNX27) is reduced in Down’s syndrome individuals (Wang et al., 2013) and it has been demonstrated that SNX27 has a role in the delivery of AMPA receptors during synaptic activity (Loo et al., 2014). Remarkably, microRNA-155 overexpression (encoded on chromosome 21) decreases SNX27 levels, which increase Aβ production by altering glutamate receptor recycling in DS (Wang et al., 2013). This study shows the molecular mechanisms underlying Aβ-dependent pathogenesis in both DS and AD. In mouse models of AD, there is a reduction in the number of AMPAR, such as in old 3xTg-AD mice (Cantanelli et al., 2014) and double knock-in (2 × KI) mice carrying human mutations in APP and presenilin-1 genes (Chang et al., 2006) as well as in human brain tissue from AD patients (Zhang et al., 2019). Soluble Aβ oligomers decrease AMPAR surface expression (Hsieh et al., 2006), by reducing the basal levels of Ser-845 phosphorylation (Miñano-Molina et al., 2011) while increasing the endocytosis of GluA2- containing AMPAR (Zhao et al., 2010).

In our protein-protein interaction network analyses, we found kalirin (KLRN) as a hub connecting with other downregulated postsynaptic proteins in DS and AD. Kalirin is localized to dendritic spines in cortical pyramidal neurons and plays a role in structural and functional plasticity at excitatory synapses. Kalirin knockdown leads to spine loss and synaptic expression of AMPA receptors and is downregulated in AD (for review see Penzes et al., 2012).

On the other hand, further GO-term-based protein enrichment analysis indicated that axon, cytoskeleton, microtubule binding, phosphorylation, and vesicle functions were affected in the upregulated protein cluster in DS and AD compared to controls. These upregulated functions suggest that the development of neuronal projections may be similarly altered in DS and AD. In the literature, the proliferation of dystrophic neurites has been described, as it occurs in AD (for review see De la Monte et al., 1999). DS-cultured neurons overexpressing APP presented increased adhesion and altered axon contact guidance (Sosa et al., 2014). In our protein-protein interaction network analyses, we found upregulation of neural cell adhesion molecule 1 (NCAM1) and 2 (NCAM2) as a hub connecting with other upregulated postsynaptic proteins in DS and AD. NCAM is implicated in cell-cell adhesion, neurite outgrowth, and synaptic plasticity. Notably, NCAM has been established as a biomarker for AD because the plasma of AD subjects shows high levels of NCAM (Chen et al., 2018) as well as in dementia of the Alzheimer type (DAT) (Todaro et al., 2004). Moreover, NCAM2 is located on chromosome 21 and is potentially involved in DS (Paoloni-Giacobino et al., 1997).

There is also evidence of microtubule alterations in our upregulated protein clusters in DS and AD patients. As broadly described in literature, microtubule alterations have been described to participate in DS and AD pathologies. Both pathologies present with hyperphosphorylation of Tau protein, which accumulates as neurofibrillary tangles (NFTs) (Ryoo et al., 2007). Remarkably, in our samples, we found upregulation of microtubule-associated protein tau (MAPT) as a hub connecting with other upregulated postsynaptic proteins involved in Ds and AD. Our results are in line with the role of microtubule-associated protein tau (MAPT) in the pathophysiology of Down syndrome and Alzheimer’s disease.

In addition, vesicle functions such as synaptic vesicle endocytosis, clathrin endocytosis, and clathrin-sculpted GABA transport vesicles were enriched in upregulated protein clusters in DS and AD. In cortical pyramidal neurons of AD and DS subjects, endocytic uptake and recycling are activated, and early endosomes are enlarged (Cataldo et al., 2000). In our samples, we found upregulation of synaptotagmin 1 (SYT1) as a hub connecting with other upregulated postsynaptic proteins involved in Ds and AD. One explanation for this result may be that postsynaptic samples are enriched by applying the subfractioning protocol, but the protein isolation is not complete, and some presynaptic proteins, such as SYT1, may be present in the samples. However, another explanation could be that some studies have demonstrated that SYT1 may also play a role in the postsynaptic trafficking of receptors. For example, SYT1 and SYT7 postsynaptic synaptotagmins mediate AMPA receptor exocytosis during LTP in CA1 hippocampal pyramidal mouse neurons and (Wu et al., 2017). In agreement with the downregulation of AMPA receptors in postsynaptic fractions of DS and AD that we found in this study, the postsynaptic upregulation of SYT1 may lead to an increased exocytosis of AMPAR as a common mechanism of DS and AD pathologies. Additionally, studies have characterized SYT1 as a novel biomarker for AD because CSF levels of synaptotagmin-1 are significantly increased in AD patients (Örhfelt et al., 2016).

Proteins such as Glutamate Decarboxylase 1 (GAD1) and the vesicular inhibitory amino acid transporter encoded by the SLC32A1 gene, which are upregulated and involved in clathrin-sculpted GABA transport vesicles and interact with GABA_B_R, were also upregulated in the postsynaptic samples of DS and AD in our study. Different DS models have shown an increased number of neurons expressing GAD1 (Souchet et al., 2014) and in a 5xFAD AD mouse model with GAD1 haploinsufficiency decreased amyloid β production was detected (Wang et al., 2017a). Additionally, a GABA_B_R antagonist restored LTP in acute slices of the dentate gyrus (DG) Ts65Dn mouse (Kleschevnikov et al., 2004; Kleschevnikov et al., 2012) suggesting a possible upregulation of GABA_B_R in the DG of Ts65Dn mice. GABA_B_R antagonists in preclinical and clinical trials have demonstrated efficacy reducing cognitive alterations in patients with mild cognitive impairment (MCI) (Froestl et al., 2004).

Additionally, these functional analyses revealed that protein kinase activity, protein ser/threo kinase activity, and phosphorylation were enriched in the upregulated cluster of proteins in DS and AD, suggesting alterations in protein kinase function and phosphorylation. In agreement with our data, previous work of Fernandez and collaborators showed an increased amount of phosphopeptides in crude synaptosomes of Ts65Dn mice (Fernandez et al., 2009). In addition, previous data from the laboratory demonstrated an upregulation of kinases in hippocampal postsynaptic samples of Ts65Dn mice (Gómez de Salazar et al., 2018). Aberrant phosphorylation has been extensively studied in AD as a critical process in AD pathology (for review see Perluigi et al., 2016) These changes in protein phosphorylation induce Aβ accumulation (for review see Oliveira et al., 2017). As already explained, Dyrk1A is overexpressed in individuals with DS and AD (Ferrer et al., 2005; Kimura et al., 2007; Park et al., 2009) and phosphorylates tau. Importantly, Tau is phosphorylated at many different sites via several protein kinases and at Ser/Thr residues in Ser/Thr-Pro sequences (for review see Kimura et al., 2014), a function that we have found to be upregulated in DS and AD samples. Through the well-described Tau hyperphosphorylation in DS and AD (Ryoo et al., 2007), some other kinases potentially dysregulated in DS and AD have been implicated in both pathologies, such as Src kinase Fyn, which also plays a role in Aβ toxicity (Ittner et al., 2010), or glycogen synthasekinase-3β(GSK-3β) (Perluigi et al., 2014). However, as a limitation of our data, we did not find conclusive upregulated levels of these kinases with a role in tau hyperphosphorylation. We did not find upregulation of Dyrk1A in the postsynaptic samples from DS and AD. In our previous proteomic study, we were unable to detect DyrK1a upregulation in the hippocampal postsynaptic fractions (Gómez de Salazar et al., 2018). This may be explained by the presence of Dyrk1a at different subcellular locations in the nucleus, cytosol, and synaptic membranes (Martí et al., 2003; Aranda et al., 2008). In our previous work, we found a significant decrease in postsynaptic Fyn kinase levels and a concomitant increase in the extrasynaptic fraction of Ts65Dn hippocampal samples. This suggests that in a pathological context, a kinase pool can be upregulated close to the synapse, in the extrasynaptic fraction and cytosol, and in hyperphosphorylated proteins after synaptic activity induction. Therefore, this upregulation may not be observed in the postsynaptic fraction under basal conditions, as in our study.

Further studies are required to better understand the mechanisms underlying DS and AD. Owing to the technical complexity of the subfractioning isolation protocol, the number of samples is limited. Increasing the number of replicates of our proteomic study may provide insight into the probable common mechanisms that we found to be inconclusive, but that have already been described in the literature, at least in one of both pathologies. Moreover, a comparison between hippocampal Ts65Dn postsynaptic samples and hippocampal AD postmortem tissue samples with cortical Ts65Dn postsynaptic samples and entorhinal AD postmortem tissue samples from this study could provide information on how different brain areas are affected by DS and AD. Complementary biochemical experiments validated the altered common mechanisms found in proteomic analyses.

In summary, our study comparing the Ts65Dn animal model and the entorhinal cortex of AD subjects revealed a possible common altered postsynaptic composition of the adult Ts65Dn mouse cortex and entorhinal cortex of AD subjects. These postsynaptic protein alterations might functionally underlie Ts65Dn-associated and AD-associated dendritic deterioration, microtubule organization, glutamatergic transmission, and phosphorylation alterations. Importantly, although further extensive mechanistic studies remain elusive, these commonly altered protein clusters in DS and AD can provide insight into the common pathological synaptic mechanisms of both diseases and, in particular, the earliest stages of AD disease progression. Moreover, our proteomic data suggest potential drug targets for modulating the commonly altered postsynaptic mechanisms in DS and AD.

## Acknowledgments

The author thanks Prof. Xavier Altafaj for providing murine samples, materials, and laboratory space at IDIBELL. Additionally, Bellvitge Biobanc was used for postmortem human samples.

